# Specific sequence of arrival promotes coexistence via spatial niche preemption by the weak competitor

**DOI:** 10.1101/2021.10.06.463344

**Authors:** Inês Fragata, Raul Costa-Pereira, Mariya Kozak, Agnieszka Majer, Oscar Godoy, Sara Magalhães

## Abstract

Historical contingency, such as the order of species arrival, can modify competitive outcomes via niche modification or preemption. However, how these mechanisms ultimately modify stabilising niche and average fitness differences remains largely unknown. By experimentally assembling two congeneric spider mite species feeding on tomato plants during two generations, we show that order of arrival affects species’ competitive ability and changes the outcome of competition. Contrary to expectations, order of arrival did not cause positive frequency dependent priority effects. Instead, coexistence was predicted when the inferior competitor (*Tetranychus urticae*) arrived first. In that case, *T. urticae* colonised the preferred feeding stratum (leaves) of *T. evansi* leading to spatial niche preemption, which equalised fitness and reduced niche differences, driving community assembly to a close-to-neutrality scenario. Our study demonstrates how the order of species arrival and the spatial context of competitive interactions can jointly determine whether species can coexist.

## Introduction

Priority effects are broadly defined as the process by which historical contingencies in community assembly (e.g. order and/or timing of arrival) change the outcome of interspecific interactions (Chase 2003; Fukami 2015). Inhibitory priority effects, when earlier arrival by one species inhibits the growth of the species arriving next, are expected to result in alternative stable states hampering coexistence (Chase 2003; Fukami 2015; Ke & Letten 2018). In turn, facilitative priority effects, when population growth is higher if individuals arrive after the settlement of a first species, do not always promote coexistence. Rather, the outcome depends on the interaction strength among species and on the environmental context in which they interact (Bulleri *et al*. 2016; Bimler *et al*. 2018). These effects have been less often observed in natural communities (Queijeiro-Bolaños *et al*. 2017; Clay *et al*. 2019; Halliday *et al*. 2020). Two major mechanisms are predicted to cause priority effects: niche preemption, in which early colonisers reduce the amount of resource available to late colonisers, and niche modification, in which the species arriving first modifies the environment, thereby inhibiting or facilitating later colonisation (Kardol *et al*. 2013; Vannette & Fukami 2014; Fukami 2015; Delory *et al*. 2019, 2021; Grainger *et al*. 2019). Niche preemption in plant communities was found to be strong in environments with high nutrient supply, as early arriving plants grew quickly and prevented growth of later colonisers by depleting space and light (Kardol *et al*. 2013). Niche modification was also detected in plants, as early colonisations modified the soil metabolome and inhibited population growth of forb, but not grass species arriving later (Delory *et al*. 2021). Although distinguishing among niche preemption and modification is not always possible (Grainger *et al*. 2018; Boyle *et al*. 2021), recent advances in coexistence theory can serve as a powerful approach to better understand the importance of historical contingencies for species coexistence. Yet the combination of these theoretical tools has seldom been applied in empirical settings.

Modern coexistence theory posits that the long-term persistence of competing species (i.e., species coexistence) can be attained by two non-mutually exclusive mechanisms: (i) equalising mechanisms that reduce average fitness differences, and therefore, dominance between species and (ii) stabilising mechanisms, which stabilise the interaction between competitors by increasing the strength of intraspecific competition relative to interspecific competition (Chesson 2000). Therefore, species will stably coexist if stabilising niche differences are larger than differences in fitness between competitors. Otherwise the species with higher fitness will eventually dominate the community (Chesson 2000; Barabás *et al*. 2018; Spaak & De Laender 2021). Under this framework, priority effects are strictly defined as positive frequency dependence (i.e., via negative niche differences), leading to the dominance of the early-arriving species (Ke & Letten 2018; Grainger *et al*. 2019; Spaak & De Laender 2021). Hence, species cannot coexist unless there is spatial variability in the order of arrival. Although recent theory offers predictions on the outcome of coexistence in systems with historical contingencies, empirical tests are conspicuously lacking (but see Cardinaux *et al*. 2018; Grainger *et al*. 2019; Song *et al*. 2020). Therefore, there is as yet scarce knowledge of which species traits interact with historical contingencies to determine outcomes of interspecific interactions.

For herbivore communities, habitat use and dispersal capacity can affect resource use and ultimately the spatial distribution of consumers. This may lead to niche preemption, as herbivores generally have preferred plant strata and the first arriving species may monopolise that resource (Grainger *et al*. 2018; Godinho *et al*. 2020a). Moreover, herbivores often induce defences on the plants they colonise, which is expected to entail niche modification for species arriving later (Erb *et al*. 2011; Moreira *et al*. 2015; Stam *et al*. 2017). For example, Hougen-Eitzman & Karban (1995) showed that early colonisation of grape vine leaves by Willamette mites negatively affected the growth of Pacific mites, probably due to systemic induction of defences. Other herbivore species can instead down-regulate plant defences, improving the performance of later colonisers (Sarmento *et al*. 2011a; Godinho *et al*. 2016), thereby potentially causing facilitative priority effects. Overall, given the environmental heterogeneity that herbivores experience (e.g., variation in leaf quality within and between plants), effects of the order of arrival on species coexistence are expected to be prevalent in these systems (Utsumi *et al*. 2010; Erb *et al*. 2011; Moreira *et al*. 2015; Stam *et al*. 2017, 2018; Godinho *et al*. 2020a). Still, what type of competitive outcome we should expect is unclear. Indeed, although the order of arrival is linked to priority effects, the interaction between the chronology of community assembly and the impact of species on the environment (e.g. where they growth and how they modify the habitat) can result in diverse outcomes, from competitive exclusion to species coexistence. Applying modern coexistence theory to this open question can shed light on the proximate mechanisms that allow for species to coexist under varied historical contingencies.

Here, we investigate the drivers of competitive outcomes by combining theoretical and empirical tools to test the mechanisms through which order of arrival affects species coexistence. We use as a model system the two closely-related competing herbivorous species, the spider mites *Tetranychus urticae* and *T. evansi. Tetranychus evansi* generally outcompetes *T. urticae* on tomato plants (Sarmento *et al*. 2011b; Orsucci *et al*. 2017; Alzate *et al*. 2020), although both species are also commonly observed on the same location (Ferragut *et al*. 2013). Niche modification is expected to be at play in this system, because the two species interact with plant defences. Indeed, *T. evansi* suppresses plant defences (Sarmento *et al*. 2011a; Alba *et al*. 2014), whereas most *T. urticae* populations induce them (Kant *et al*. 2008). This asymmetrical niche modification is predicted to increase the probability of coexistence by hampering growth of the stronger competitor and favouring growth of the inferior one, when the later arrives on plants colonised by the other species. Moreover, niche preemption may occur, as both *T. evansi* and *T. urticae* prefer the upper, more nutritious leaves of tomato plants, where their performance is higher (Godinho *et al*. 2020a). Thus, early-arriving species could occupy the preferred niche and displace the other species to lower, less optimal, plant strata. We tested this by performing a series of multi-generational experiments where we varied order of arrival and measured space use by the two competing species. To quantify the magnitude of niche modification, we tested how these species modify the expression of genes associated with induced defenses on tomato. We then applied modern coexistence theory to unravel the conditions favouring coexistence or potentially leading to priority effects.

## Material and Methods

### Model system, species characteristics, and maintenance of experimental populations

*Tetranychus urticae* is a generalist herbivore that feeds on many economically important crops (Helle & Sabelis 1985; Grbić *et al*. 2011; Sousa *et al*. 2019), whereas *T. evansi* is a solanaceous specialist that has recently invaded Europe (Boubou *et al*. 2012). Both species colonise tomato plants, although *T. urticae* may shift to other hosts if *T. evansi* is present (Ferragut *et al*. 2013).

All experiments were performed with outbred populations of *T. urticae* and *T. evansi* spider mites, formed via controlled crosses among four *T. evansi* and three *T. urticae* populations collected in different locations in Portugal (Godinho *et al*. 2020b). Populations were maintained in boxes containing leaves detached from five-week-old tomato plants (*Solanum lycopersicum, var* MoneyMaker), with their petiole in a small pot containing water Twice a week, overexploited leaves were removed, and water and new tomato leaves were added. Before infestation, tomato plants were kept in a separate climatic chamber and watered three times per week. Mites and plants were kept under controlled conditions (25 °C, 70% humidity, 16 /8 L/D hours).

We created same-age cohorts of mated *T. urticae* and *T. evansi* females for each block. To this aim, females were placed during 48h in petri dishes (14.5 cm diameter, with a layer of wet cotton watered twice per week) and two freshly cut tomato leaves. One week later, another tomato leaf was added. In the experiment, we used females with 13-15 days of age.

### Theoretical approach for predicting competitive outcomes: quantifying niche and fitness differences

Data collected in the experiments were used to parameterise a mathematical model from which niche and average fitness differences can be quantified to then draw predictions of competitive outcomes. We assume that the population dynamics in our experiment can be described by a Beverton-Holt function (Levine & HilleRisLambers 2009; Godoy & Levine 2014):

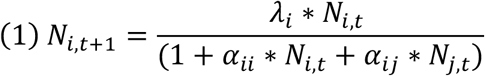

Where *N*_*i,t+1*_ is the number of individuals of species *i* in the next generation, *λ*_*i*_ the intrinsic growth rate of species *i* in absence of competitors, *α*_*ii*_ the intraspecific competitive interaction describing the per-capita effect of species *i* on itself, *α*_*ij*_ the interspecific competitive interactions describing the per-capita effect of species *j* on species *i*, and *N*_*i,t*_, *N*_*j,t*_ the number of individuals of species *i* and *j* in the current generation, respectively. We assume that spider mites do not have a dormant stage. Thus, *λ*_*i*_ represents the fraction of eggs that hatch and become females that reproduce in the next generation. One of the predictions of modern coexistence theory is that, for species to coexist, they must invade the resident species from rare. Because for our system equilibrium densities are difficult to attain within a time frame fast enough to study the impact of priority effects on species coexistence, we instead used experimental gradients of density and relative frequency to estimate intra and interspecific competitive interactions (the α’s) and intrinsic growth rate (λ) for each species, an approach well established and validated by previous work (Godoy & Levine 2014; Matías *et al*. 2018; Song *et al*. 2020).

From the above mentioned model, niche overlap (*ρ*) is defined as follows (see details in Chesson 2012; Godoy & Levine 2014).

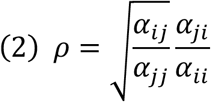

This formula reflects the average degree to which species limit individuals of their own species relative to heterospecific competitors. If species limit population growth of their own species more strongly than that of their competitors (*α*_*jj*_, *α*_*ii*_, are much greater than *α*_*ij*_, *α*_*ji*_), then niche overlap will be low, favouring coexistence. Alternatively, niche overlap will approach one, which hampers stable coexistence. Stabilising niche differences are thus expressed as 1-*ρ*.

Average fitness differences 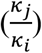 (Chesson 2012; Godoy & Levine 2014) are defined as:

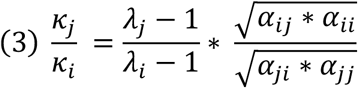

The greater the ratio, 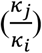, the greater the fitness advantage of species *j* over *i*. If this ratio is one, species are equivalent competitors. Coexistence requires both species to invade when rare (Chesson 2012), which is satisfied when (Godoy & Levine 2014):

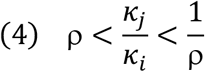

Stable coexistence is possible whenever species have either large stabilising niche differences (corresponding to small niche overlap) that overcome large average fitness differences, or at the other extreme, via an a close-to-neutral scenario (Scheffer *et al*. 2018), where, even with weak niche differences, small fitness differences stabilise the interaction between competitors. If no coexistence is predicted, we can pinpoint if this is due to competitive exclusion (when fitness differences are larger than niche differences) or to priority effects, leading to alternative states when niche differences are negative. Negative niche differences imply that each species limits the growth of the competitor more than their own (Fukami & Nakajima 2011; Ke & Letten 2018).

We used maximum likelihood techniques to parameterise the population model following a nested approach. That is, we first created a single model for which we estimate the intrinsic growth rate in absence of competitors (*λ*), and then we used this information as prior for subsequent more complex models that include intra and interspecific competitive interactions (the *α*’s) (Matías et al. 2018). *λ* values were considered fixed per species across empirical treatments, but competition varied across treatments because mite species can differentially disperse and modify leaf quality and availability (see the full details in the Supplementary Material and Methods).

### Experiments

To test the impact of order of arrival on coexistence, we performed a series of experiments in which we either manipulated the order of arrival and relative frequency (i.e., relative initial abundance with a constant density of 20 individuals), or the initial density of each of two species of competing spider mites. Furthermore, to estimate the effect of order of arrival on promoting niche preemption, we quantified leaf occupation for both species at the end of the experiment. Finally, to estimate the effect of order of arrival on promoting niche modification, we quantified induction of plant defences of both species.

In the first experiment, both species were introduced simultaneously using the following proportions of *T. evansi* : *T. urticae*: 1:19; 10:10 and 19:1, along with the single-species controls (20:0 and 0:20). To manipulate the order of arrival, we introduced (i)10 *T. evansi* females 48h before 10 *T. urticae* females and vice versa and (ii) 19 *T. evansi* females 48h before 1 *T. urticae* female and vice versa (Figure S1). The experiment was done in two blocks, one week apart. Each block contained five boxes of each experimental treatment (nine treatments, n=10), each with a pot filled with water and two freshly cut tomato leaves from five-week-old tomato plants. Leaf pairs consisted of leaves 2 and 4 or 3 and 5 (leaf number is inversely proportional to leaf age), to ensure that each box contained a younger and an older leaf, since both species prefer younger leaves (Godinho *et al*. 2020a). Adult females were distributed by the two leaves, following the treatments described above. After one generation (circa 14 days), two more leaves were added to ensure enough resources for the second mite generation. Boxes that initially received the leaf pair 2-4, received leaves 3-5 and vice versa. After two generations, we counted the number of adult females of each species on each leaf.

Next, we estimated the growth rate of each species by counting the number of adult females obtained from the progeny of a single *T. urticae* or *T. evansi* female ovipositing for 48h in two overlapping 18mm leaf disks (n=10). These disks were placed in square petri dishes with a layer of wet cotton and were watered every two days. The number of adult females produced was assessed after one generation.

#### Quantification of niche modification

To quantify the magnitude of niche modification induced by *T*.*urticae* and *T*.*evansi*, we investigated how these two species modified the expression of genes associated with plant defences. As controls, we quantified the expression of the same genes upon infestation with spider mites from *T. urticae* Santpoort and *T*.*evansi* Viçosa populations, known to induce and suppress tomato defences, respectively (Alba *et al*. 2014). Details of this experiment are given in the Supplementary Material and Methods and Table S1.

### Data Analyses

#### Effect of order of arrival and initial frequency on species abundance

To test the impact of order of arrival, frequency and their interaction on the proportion of adult females of each species after two generations, we performed the following general linear mixed model (lme4 package, Bates et al. 2015), using the binomial family:

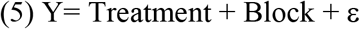

Where Y corresponds to the combination of two vectors with the number of *T. evansi* and *T. urticae* females after two generations, Treatment (fixed factor) to the combination of different orders of arrival and initial frequencies, Block (random factor) to whether the experiment was performed on week one or two, and ε to the residual error. We then performed *a priori* contrasts, using testInteractions from phia package (Rosario-Martinez 2015) as our experimental design was not orthogonal. To compare the effect of different orders of arrival, we performed contrasts between the treatments with same initial frequency but different orders of arrival. To compare the effect of frequency, we performed contrasts between treatments with same order of arrival but different initial frequencies. Contrasts were corrected for multiple comparisons using FDR correction (Benjamini & Yekutieli 2001). To test whether the results were biased by the order in which the leaf pairs were added to the boxes, we repeated these statistical analyses separately for each leaf pair.

#### Effect of order or arrival and initial frequency on leaf occupancy and aggregation

To test if coexistence outcomes could be explained by niche preemption, we compared occupancy patterns of each species across the four leaves. For the single species treatment, we tested if the number of females differed across leaves (model 6). For the double species treatment, we tested if the order of arrival, initial frequency, or their interaction changed mite distribution (model 7), by comparing it to the distribution of the single species treatment.

We applied the following binomial models, with Leaf and/or Treatment and their interaction as fixed factors, for the control (model 6) and experimental (model 7) treatments:

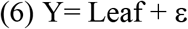

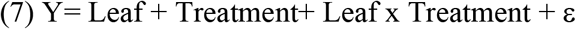

where Y corresponds to the combination of two vectors with the number of *T. evansi* (or *T. urticae)* females on each leaf and the total number of individuals on each box that were not on that leaf. To test whether the results were biased by the order in which the leaf pairs were added to the boxes, we repeated these statistical analyses accounting for the preference of each species for each leaf pair. For the double treatment, a posteriori contrasts were done between each treatment and the corresponding single species treatment. The initial fitting with Block as a random factor, indicated no variance in this factor, thus we fitted only fixed factors. We also tested in changes in order of arrival affected aggregation scores (see details in Supplementary Material and Methods).

All analyses were done using R (R Core Team 2021). To predict coexistence outcomes we used the package “cxr” (García-Callejas *et al*. 2020). Plots were done using “ggplot2” (Wickham 2016) and “cowplot” (Wilke 2020) packages. Data and scripts are available in the github repository: https://github.com/irfragata/order_arrival_niche_preemption.

## Results

### Effect of order of arrival and initial frequency on species abundance

The number of individuals of each species on tomato plants were affected by the order of arrival (contrasts between *T. evansi* arriving first vs. simultaneously: χ^2^ = 44.252, df = 1, p-value < 0.0001; or *T. urticae* arriving first vs. simultaneously: χ^2^ = 375.860, df = 1, p-value < 0.0001), and their initial frequency (contrasts between *T. evansi* starting at equal *vs*. higher frequency. : χ^2^ = 784.335, df = 1, p-value < 0.0001; or *T. urticae* starting at equal *vs*. higher frequency: χ^2^ = 654.903, df = 1, p-value < 0.0001). Specifically, the abundance of *T. evansi* females after two generations was higher when this species arrived first or simultaneously with *T. urticae*, independently of initial frequencies. However, the additional advantage provided by arriving first was much larger in the equal frequency treatment (Table S2, Fig.1). The abundance of *T. urticae* after two generations was also affected by initial frequency and order of arrival. Indeed, the final number of *T. urticae* females was higher when this species arrived first and was at high initial frequency, than in the equal frequency treatment (Table S2, Fig. 1). We observed the same patterns when performing these analyses per leaf pair (Table S3). Overall, these results confirm that *T. evansi* is a superior competitor as observed in previous studies (Sarmento *et al*. 2011b; Ferragut *et al*. 2013; Alzate *et al*. 2020).

**Figure 1.**
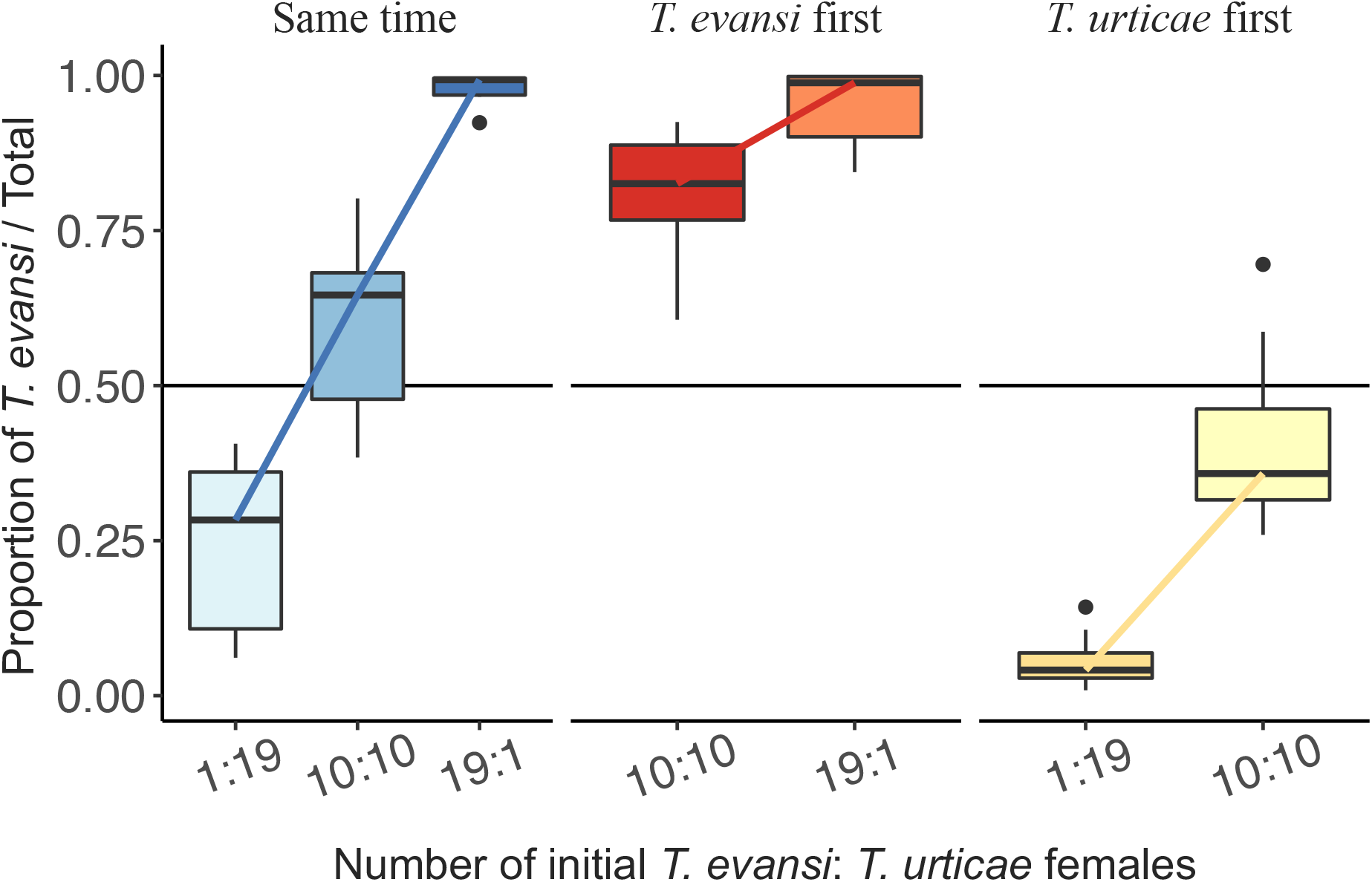
Proportion of spider mites *Tetranychus evansi* females (y-axis) depending on initial frequency (number of initial females *T. evansi*: *T. urticae*, x-axis) and order of arrival (same time vs. *T. evansi* or *T. urticae* arriving 48h before its competitor) after two generations. *Tetranychus evansi* is the better competitor overall (ratio above 0.5), unless *T. urticae* arrives first or is at higher initial frequency. A posteriori contrasts show a strong effect of order of arrival in the proportion of females of the two species (Table S2B). Initial frequency also impacts the final ratio, with a stronger effect when *T. urticae* arrives first or at the same time than *T. evansi* (Table S2B). Boxplots represent median and quartiles of the 10 boxes within treatment.

### Effect of order of arrival on coexistence

The order of arrival modified the outcome of competition between the two species. *Tetranychus evansi* (the superior competitor) is predicted to exclude *T. urticae* when it arrives first or at the same time. Under this exclusion scenario, the rate of competitive exclusion is expected to be faster when *T. evansi* arrives first due to a decrease in niche differences (Fig 2). The small overlap between the lower confidence interval with the priority effects region suggests that positive frequency dependence might also emerge in this system. Interestingly, coexistence was only possible when *T. urticae* arrived first (Fig. 2). This outcome was due to small niche and fitness differences among competitors, leading to a quasi-neutral scenario. Specifically, when *T. urticae* arrived first, we observed similar strengths of intra- and interspecific interactions among species (Fig S2A). Contrary to expectations and previous studies, the order of arrival was not associated with positive frequency dependence leading to priority effects. However, since the order of arrival modified the outcome of the interactions between *T. urticae* and *T. evansi*, we can also interpret these results as priority effects (*sensu* (Chase 2003; Fukami 2015) allowing for coexistence between species in our system.

**Figure 2.**
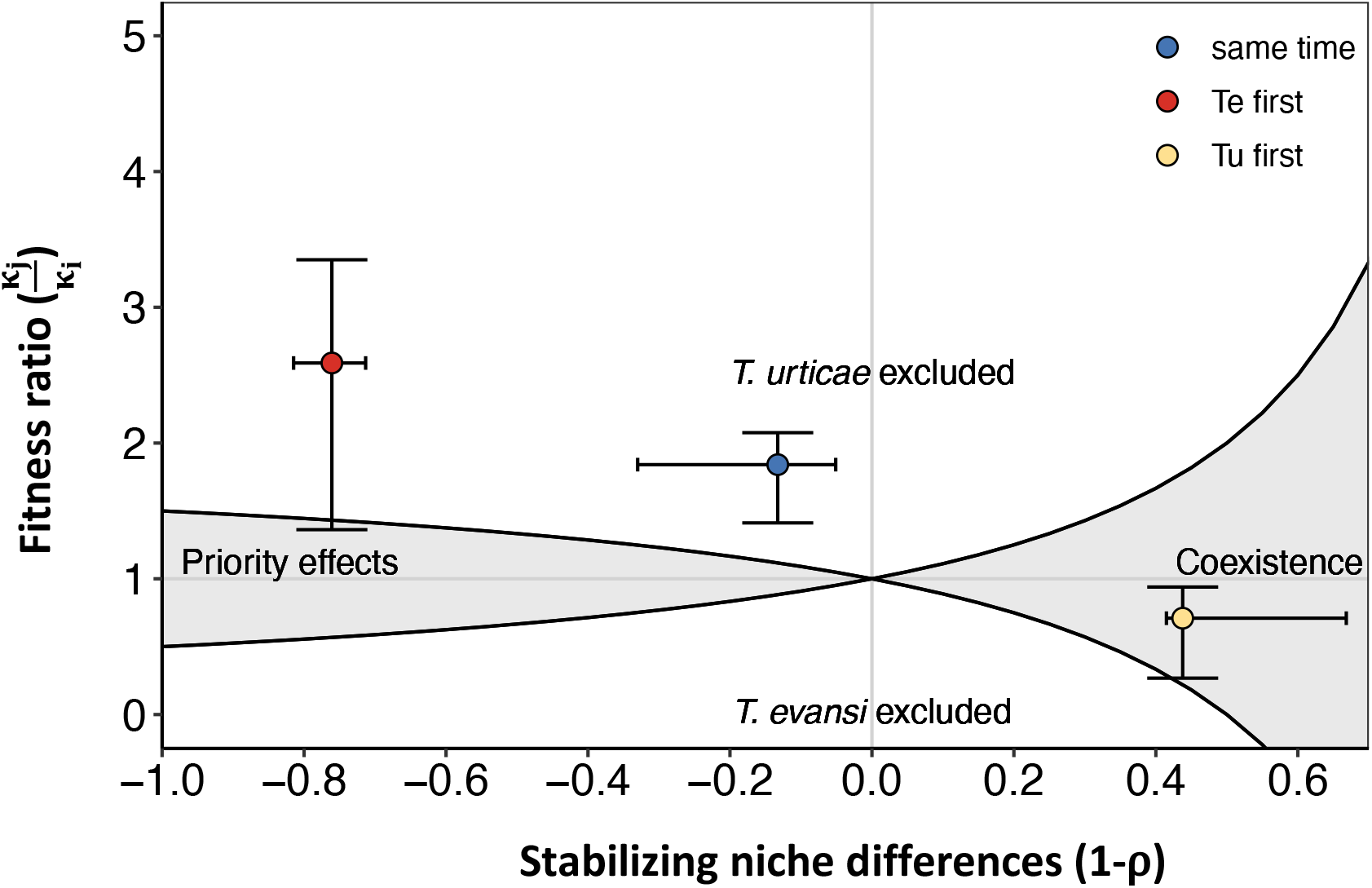
Relationship between average fitness differences 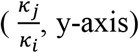 and stabilising niche differences (1-ρ, x-axis) for different orders of arrival (*Tetranychus evansi* first – red, same time – blue, *T. urticae* first – yellow). Plotting average fitness differences against niche differences allows mapping different competitive outcomes predicted by modern coexistence theory. The coexistence condition (eq. 4) and its symmetrical for each competing species, represented by the two solid black lines, allow defining the space in which species can coexist due to negative frequency dependence or enter alternative stable states due to positive frequency dependence, whenever niche differences are greater or smaller than zero respectively. Otherwise, the species with higher fitness will exclude the other. In our case, the only scenario in which species are predicted to coexist is when *T. urticae* arrives first (yellow). Error bars for each outcome indicate the 95% confidence interval from the maximum likelihood estimates. For the other two cases, it is predicted that the superior competitor *T. evansi* will exclude *T. urticae*.

### Effect of order of arrival and initial frequency on leaf occupancy and aggregation

When *T. evansi* was alone, it reached higher abundances on leaves 3 and 4 (Table S4A, Fig 3B), whereas *T. urticae* was less abundant on leaf 2 in comparison to all others (Table S4A, Fig 3D). Fewer *T. evansi* females were found on leaf 4 when *T. urticae* arrived first, and on leaf 3 when *T. urticae* started with higher frequency and both species arrived at the same time (Fig 3, Fig S3A, Table S4B). When *T. evansi* arrived first or started at higher frequency, we observed fewer changes on its own leaf occupancy (Fig S4A). The distribution of *T. urticae* showed a slight shift when it arrived first, with a reduction on the prevalence of leaves 2 and 5 and slightly higher occupation of leaves 3 and 4 (Fig. S3B, Table S4B). When *T. evansi* started at high frequency, there was also a shift in *T. urticae* distribution, with a lower occupancy of leaves 2 and 5 (Fig S3B). We observed similar shifts in leaf occupation when performing the analyses accounting for the order in which each leaf pair was added (Fig S4, Table S5).

**Figure 3.**
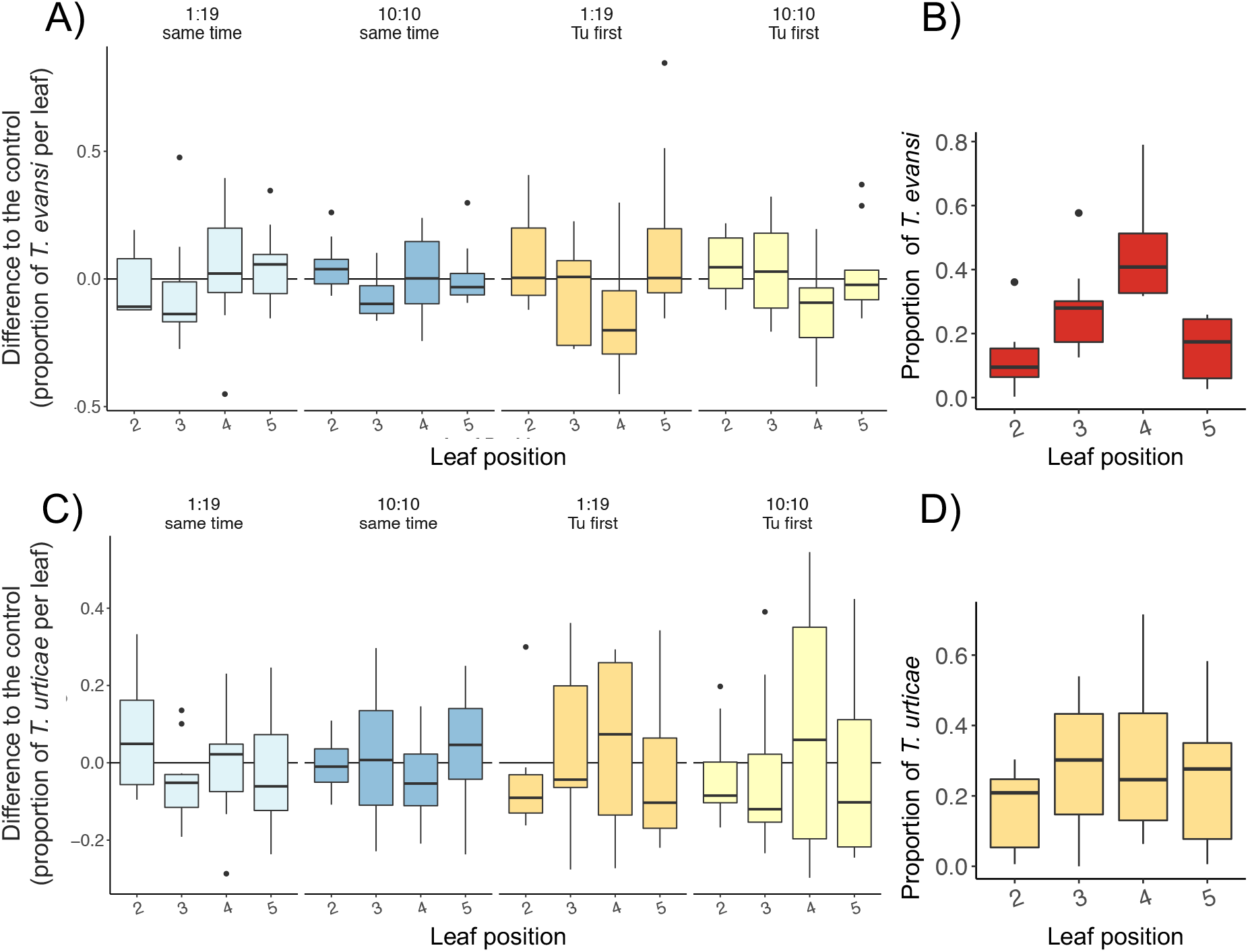
Differences between expected and observed leaf occupancy for *Tetranychus evansi* (A) and *T. urticae* (C) for a subset of the experimental treatments (when *T. urticae* arrived first or at the same time as *T. evansi*, note that Figure S3 includes all treatments); leaf occupancy for *T. evansi* (B) and *T. urticae* (D) in the control, single species, treatments. Leaf 2 corresponds to the oldest leaf and leaf 5 to the youngest. For each box, we calculated for each species the proportion of females occupying each leaf in relation to the total number of females present. For the experimental treatments we calculated the difference between this proportion and the average proportion for the control treatments. Thus, positive values indicate that there are more females on that leaf than expected based on the single-species treatment and negative values indicate the reverse pattern. Overall, we see that. *T. evansi* reduces occupancy on leaf 4 when *T. urticae* arrives first and on leaf 3 when the two species arrive at the same time. In contrasts, *T. urticae* shows a slight increase in occupancy of leaf 4 when it arrives first and a slight decrease in occupancy of leaves 2 and 5.

Spatial aggregation significantly differed among treatments (χ^2^ = 18.186, df = 6, p-value = 0.01279), being higher in treatments with similar initial densities (cf. Fig S5 with Fig 1, Table S5). We observed a significant difference in C-score, with higher aggregation when both species arrived at the same time and had equal frequency, and a lower aggregation when both species arrived at the same time and *T. evansi* started at higher frequency (Table S6). Order of arrival did not change the C-score (Fig. S5, Table S6).

### Quantification of niche modification

Plants infested by *T. urticae* or *T. evansi* populations showed patterns of gene expression similar to those of Viçosa, the suppression control, and significant differences with Santpoort, the induction control (Fig. S6; Table S7). We thus conclude that, both populations suppress plant defences.

## Discussion

This study shows that order of arrival interacts with competitive ability to determine the probability of coexistence between congeneric species that share common resources. When both species arrive at the same time or the superior competitor (*Tetranychus evansi*) arrived first, *T. urticae* was predicted to be excluded. Coexistence was only predicted when the inferior competitor (*T. urticae*) was the first species colonising the habitat. Analyses of leaf occupancy show that these competition outcomes are linked to a spatial niche preemption process in which *T. evansi* was displaced from its preferred food stratum when *T. urticae* arrived first. As a result of this complex interaction between order of arrival, species competitive ability, and spatial occupancy, we observed a particular configuration that allows species coexistence: both species equalised their fitness differences to the extent that they can coexist despite small niche differences. These multiple lines of evidence challenge the common understanding of the inhibitory role of niche preemption in coexistence between species.

We found that *T. evansi* had higher competitive ability and growth rate, and often excluded *T. urticae* (Fig 1, 2). This is in line with laboratory observations showing that *T. evansi* outcompeted *T. urticae* on tomato plants (Sarmento *et al*. 2011b; Alzate *et al*. 2020, but see Orsucci *et al*. 2017) and with field observations showing a shift in host use in *T. urticae* upon invasion by *T. evansi* (Ferragut et al. 2013). Still, these two species can co-occur in the field in the same plant species (Ferragut *et al*. 2013; Orsucci *et al*. 2017; Zélé *et al*. 2018). The advantage created by the earlier arrival of *T. urticae*, and associated reduction in interspecific competition by *T. evansi*, could be one of the possible mechanisms fostering their coexistence. Indeed, *T. urticae* can withstand colder temperatures than *T. evansi* (Gotoh *et al*. 2010; Khodayari *et al*. 2013; Riahi *et al*. 2013; White *et al*. 2018), hence it is expected to arrive first in the season. Field surveys that sample both species in the same location across seasons are needed to further explore this hypothesis.

Historical contingencies emerging from order of arrival can happen through two main mechanisms: niche modification or niche preemption (Fukami 2015). In our system, niche modification may arise via interactions between spider mites and plant defences. However, we observe that both species suppress plant defenses. If suppression would affect species performance, we would expect higher production of offspring when the competitor arrives first. We did not observe this, suggesting that this mechanism of niche modification does not affect the outcome of competition in this system.

Niche preemption can occur through monopolisation of nutrients or space, which can be particularly important among competitors with similar requirements (Grainger *et al*. 2018; Holditch & Smith 2020). In our study, we observed a shift in the leaf occupancy pattern of *T. evansi* females when *T. urticae* arrived first. This displacement of *T. evansi* from the preferred food stratum (i.e., younger, more nutritious leaves) by early-arriving *T. urticae* can explain the decreased performance of the superior competitor. Thus, our results indicate that variation in species performance driven by habitat quality heterogeneity (Orians *et al*. 2000; Orians & Jones 2001) combines with order of arrival to generate niche preemption, providing a mechanism for the two herbivores to coexist.

Order of arrival is a major determinant of community assembly across diverse taxa, from microbes to plants (Chase 2003; Erb *et al*. 2011; Kardol *et al*. 2013; Stam *et al*. 2017; Grainger *et al*. 2018, 2019; Clay *et al*. 2019, 2020; Halliday *et al*. 2020). Most of these studies show that early colonisers inhibit growth and decrease performance of late arriving species, especially in those that occupy very similar niches (Fargione *et al*. 2003; Vannette & Fukami 2014; Delory *et al*. 2019, 2021; Grainger *et al*. 2019), although very few concern herbivorous species competing for the same niche (e.g. Grainger *et al*. 2018; Holditch & Smith 2020). Other studies found that order of arrival does not affect community assembly (e.g. Delory *et al*. 2021) or that initial colonisers facilitate later colonisation of other species (e.g. Queijeiro-Bolaños *et al*. 2017; Delory *et al*. 2019). Here, we show that coexistence is promoted by niche preemption because early colonisation by the inferior competitor leads to increased intraspecific competition for the superior competitor and reduced interspecific competition for itself.. As a result, both species can coexist under a quasi-neutral scenario because this equalising effect on fitness differences is enough to fit within the constraints of small niche differences. Our study adds a novel perspective to the growing body of evidence that historical contingencies shape ecological communities, by showing that the probability of coexistence of two competing herbivores changes due to an interaction between order of arrival and species competitive ability.

Priority effects were recently incorporated into modern coexistence theory (Ke & Letten 2018; Spaak & De Laender 2021), but empirical tests quantifying the effects of order of arrival on species coexistence remain very rare. In another study, Grainger *et al*. (2019) documented that positive frequency dependence, due to strong priority effects, arose from changes in order or arrival in yeast species feeding on floral nectars. In contrast, our results show that order of arrival did not lead to alternative states caused by priority effects under positive frequency dependence. Rather, we predicted either competitive exclusion when *T. evansi* arrived first because it excluded *T. urticae* or coexistence when *T. urticae* arrived first. Overall, these results suggest that in this system deterministic expectations, stemming from theory, can be strongly influenced by small stochastic events, such as changes in order of arrival, because it affects the timing of dispersal across and within host plants.

Framing priority effects in the modern coexistence theory (Ke & Letten 2018) is undoubtedly an important step to mechanistically understand how order of arrival affects community assembly processes. However, in this framework, priority effects are only caused by positive frequency dependence (i.e., population growth rate is higher as individuals become relatively more abundant) (Fukami 2015; Song *et al*. 2020). Including other types of interactions and outcomes into modern coexistence framework is fundamental to improve our ability to understand how species coexist (Spaak *et al*. 2021). Here we show that order of arrival can lead to coexistence via niche preemption by the inferior competitor. Thus, our results show that changes in the order of arrival can produce a wide range of competitive outcomes from coexistence to competitive exclusion due to positive and negative frequency dependence. Therefore, it is urgent that ecologists widen the scope of the multiple outcomes that historical contingency can produce on species coexistence.

Most empirical and theoretical studies emphasize the inhibitory nature of niche preemption (Fargione *et al*. 2003; Fukami 2015; Vieira *et al*. 2018; Delory *et al*. 2019), with the early arriving species outcompeting the other. However, recent theory suggests that, in a resource competition model of two species, niche preemption by the inferior competitor could facilitate coexistence under a trade-off between order of arrival and the resource levels of zero net growth (R*) (Qi et al. 2021). Our study is, to the best of our knowledge, the first empirical study showing that niche preemption by the weaker competitor promotes coexistence. This striking change in the outcome of competitive interactions emerges mostly due to a decrease in niche overlap, shifting niche differences from negative to positive. This suggests that even small differences in order of arrival can be sufficient for the monopolisation of a resources in plant-herbivore interactions, which may suffice to allow coexistence between competitor species. Therefore, our results demonstrate how small temporal differences percolate into small spatial heterogeneities, fostering coexistence and the maintenance of diversity.

## Acknowledgments

This work was financed by an ERC (European Research Council) consolidator grant COMPCON, GA 725419 attributed to SM and by FCT (Fundação para Ciência e Tecnologia) with the Junior researcher contract (CEECIND/02616/2018) attributed to IF. RC-P is supported by grant #2020/11953-2 São Paulo Research Foundation (FAPESP) and grant R-2011-37572 Instituto Serrapilheira. OG acknowledges financial support provided by the Spanish Ministry of Economy and Competitiveness (MINECO) and by the European Social Fund through the Ramón y Cajal Program (RYC-2017-23666). AM was funded by National Science Centre, Poland (grant no. 2018/28/T/NZ8/00060) and Excellence Initiative - Research University programme (support for the internationalization of the Adam Mickiewicz University PhD students, no. 003/13/UAM/0018). The authors acknowledge stimulating discussion with all members of the Mite Squad, in particular Flore Zélé and Diogo Godinho, which have significantly improved the experimental design and interpretation of the results, and Marta Artal for invaluable infrastructure at meetings in Sevilla.

