## Supplementary information for "Specific sequence of arrival promotes coexistence via spatial niche preemption by the weak competitor"

### Supplementary Material and Methods

***Quantification of gene expression in plants infested by each spider-mite species (niche modification)***

To quantify the magnitude of niche modification induced by *T.urticae* and *T.evansi*, we investigated how these two species modified the expression of genes associated with plant defenses, using five-week-old tomato plants of the Castlemart variety (developing in the same conditions as described above). As controls, we quantified the expression of the same genes upon infestation with spider mites from *T. urticae* Santpoort and *T.evansi* Viçosa populations, known to induce and suppress tomato defenses, respectively (Alba et al. 2014) (n=9 per population). We cut leaf 4 and contoured the petiole of second clockwise top leaflet with lanoline to prevent mites from dispersing to other leaflets. We then infested this leaflet with 30 adult mated females of each population. After four days, a 14mm disc was isolated from the infested leaflet, which was immediately placed in SafeLock 2mL tube and immersed in liquid nitrogen. All samples were stored at -80ºC before RNA isolation. Keeping the plant material frozen, we crushed the leaf discs using Mixer Mill (Retsch MM400) operating the equipment for 15 seconds at 30 Hz. After converting it to dust, we placed the tubes in liquid nitrogen, then added 600 uL of extraction buffer (acid (pH=4.3±0.2) phenol – 0.1M LiCl, 100 mM Tris-HCl (pH=8), 10 mM EDTA, 1% SDS [1:1]) and homogenized the mixtures by 30 seconds vortex followed by 5 minutes of rest at room temperature. Then we added 300 uL chloroform-isoamylalcohol (24:1), vortexed and centrifuged for 10 minutes at 12000 rpm. We removed the water phases, always maintaining samples on ice, and, in new tubes, mixed with one volume of 4M LiCl. Following overnight RNA precipitation at -20ºC, we placed the tubes at -80ºC for 30 minutes, defrosted and centrifuged for 30 minutes (12000 rpm) at 4ºC. We then dissolved the pellets in 250 uL of water (for molecular biology), maintaining all samples on ice, then incubated for 10 minutes at 65ºC and added 0.1 volume of 3M NaOaC (pH=5.2) and 2 volumes of cold 96% ethanol followed by 30 minutes of precipitation at -20ºC. After centrifugation for 30 minutes (12000 rpm) at 4ºC we washed the RNA pellets with 180 uL of cold 70% ethanol and centrifuged again for 5 minutes (12000 rpm) at 4ºC. Pellets were dried in SpeedVac Vacuum Concentrator (Thermo Fisher Scientific) for 3-5 minutes and finally dissolved in 40 uL of water (for molecular biology) by incubating the tubes at 65ºC for 15 minutes. The total RNA isolation protocol used was adapted from Verwoerd et al. (1989) with minor modifications. After the initial RNA isolation, two micrograms of RNA were DNAse-treated with Ambion Turbo DNA-free kit (Invitrogen) and cDNA was synthesized with RevertAid H Minus Reverse Transcriptase (Thermo Fisher Scientific). 1µL of cDNA was used as template for a 20µL quantitative reserve-transcriptase polymerase chain reaction (qRT-PCR) using the CFX96 Real-Time system (Bio-Rad) with SsoFast™ EvaGreen®Supermix (Bio-Rad). Each sample was run three times (three technical replicates) and the average from these was used in subsequent analyses. We selected marker genes that are known to respond to infestation by both these species (Sarmento at al. 2011, Alba et al., 2014): PI-IIc and PPO-D (jasmonic acid pathway-markers), and PR-1a (salicylic acid pathway-marker). The tomato Actin gene was used as reference to calculate the normalized expression. Gene identifiers, primer sequences and references are listed in Table S1. The normalized expression (NE) was calculated using the ΔCt method as described by Alba et al. (2014). To plot the relative expression, NE values were scaled to the treatment with the lowest average NE.

**Supplementary statistical analyses**

*Quantification of niche modification*

To test how the spider mite populations affected the expression of plant defences, we compared the normalized expression of genes from the salicylic and jasmonic acid pathways, with the expression of two control populations: *T. urticae* Santpoort population, which induces plant defences, and *T. evansi* Viçosa, which suppresses such defences. For that we ran the following general linear model:

$$(10)\ Y = Population + \varepsilon$$

Where Y is the normalized expression for the genes PI-IIc, PPO-D and PR-1a, and Population corresponds to a fixed factor coding for all analyzed populations (outbred populations and the two controls). We performed a priori contrasts between each outbred population and the controls for induction and suppression as described above.

**Aggregation**

Since *T. evansi* suppresses defences locally, it is expected that *T. urticae* aggregates with it (Sato *et al.* 2016). To test if aggregation changed with order of arrival or initial frequency, we calculated the Checkerboard score (C-score) (Gotelli & Rhode 2002) per replicate. The C-score quantifies species co-occurrence, measuring the extent to which they segregate or aggregate across environments (Gotelli & Rhode 2002). The bipartite package (Dormann *et al.* 2008) normalizes the C-score between 0 (no aggregation) and 1 (aggregation), allowing comparisons between treatments. To calculate the C-score per leaf, we created a presence-absence matrix per leaflet and leaf for each box. We then applied the following general linear mixed model to test for differences in aggregation between treatments:

$$(9)\ Y = Treatment + \varepsilon$$

Where Y is the computed C-score and Treatment is a fixed factor. Contrasts were performed between initial frequency and order of arrival, applying FDR correction for multiple comparisons, as described above. The initial fitting with Block as a random factor, indicated no variance in this factor, thus we fitted only fixed factors.

**Increasing model complexity to estimate niche and fitness differences.**

To parameterize population model with information from detailed experiments, we used maximum likelihood techniques with increasing model complexity. The underlying aim to perform this framework is to allow model convergence within realistic estimates, and without the need to constrain the parameter range. Doing so it allow us to compute in order to estimate normal errors distribution of each parameter from the Hessian matrix. Specifically, during the parameterization we used three different models in which estimates obtained from model 6A were used as priors for model 6B, and those obtained from model 6B were used as priors for model 6C.

$$\text{(6A) } N_{t+1} = \lambda * N_t$$

$$\text{(6B) } N_{i,t+1} = \frac{\lambda * N_t}{(1+\alpha * N_t)}$$

$$\text{(6C) } N_{i,t+1} = \frac{\lambda_i * N_{i,t}}{(1+\alpha_{ii} * N_{i,t} + \alpha_{ij} * N_{j,t})}$$

The initial model (6A) considers only the intrinsic growth rate in the absence of interactions ($\lambda$). This model was parameterized using estimates from the experiment with single *T. urticae* or *T. evansi* female. The intrinsic growth rate in the absence of interactions was considered constant (i.e. fixed value) in the subsequent models. Model 6B adds a parameter $\alpha$, which accounts for the overall effect of competition, and finally, model 6C separates this into the intra and interspecific components. with this procedure, we allowed intra and interspecific competition to vary across treatments to account for the fact that herbivores can modify their response to competition by dispersal events across leaves, and by modifying leaf quality and availability.

#### Supplementary Tables and Figures

Supplementary Table S1 – Information (name, gene identifier, primers used and references from where the primers were obtained) about genes used for plant defense induction/suppression quantification. These genes are widely used to test plant defence responses to infestation by *T. evansi* and *T. urticae*.

| Target Gene | Name | Gene Identifier | Forward Primer (5'→ 3') | Forward Primer (5'→ 3') | References |
|---|---|---|---|---|---|
| **PI-IIc** | Proteinase Inhibitor IIc | Solyc03g020050.2 | CAGGATGTACGACGTGTTGC | GAGTTTGCAACCCTCTCCTG | Gadea et al. (1996) |
| **PPO-D** | Polyohenol-oxidase-D | Solyc08g074680.2 | GCCCAATGGAGCCATATC | ACATTCGATCCACATTGCTG | Newman et al. (1993) |
| **PR-1a** | Pathogenesis-related protein 1a | Solyc09g007010.1 | TGGTGGTTCATTTCTTGCAACTAC | ATCAATCCGATCCACTTATCATTTTA | Van Kan et al. (1992) |
| **Actin** | Actin | Solyc03g078400.2 | TCAGCACATTCCAGCAGATGT | AACAGACAGGACACTCGCACT | Tomato Genome Consortium (2012) |

Supplementary Table S2– *A priori* contrasts to test for the impact of initial density and order of arrival on the proportion of females of the two species after two generations of contact.

| **Test** | **Contrasts** | **DF** | **Chisq** | **P-Value** |
|---|---|---|---|---|
| Order of arrival | 10:10 same time – 10:10 T. urticae first | 1 | 80.4130 | < 0.0001 |
| | 10:10 same time - 10:10 T. evansi first | 1 | 73.9895 | < 0.0001 |
| | 19:1 same time- 19:1 T. evansi first | 1 | 0.1776 | 0.6735 |
| | 1:19 same time- 1:19 T. urticae first | 1 | 298.7552 | < 0.0001 |
| | 10:10 T. evansi first – 10:10 T. urticae first | 1 | 298.8019 | < 0.0001 |
| Initial density | 10:10 same time – 19:1 same time | 1 | 213.395 | < 0.0001 |
| | 10:10 same time – 1:19 same time | 1 | 165.891 | < 0.0001 |
| | 10:10 T. urticae first – 1:19 T. urticae first | 1 | 636.484 | < 0.0001 |
| | 10:10 T. evansi first – 19:1 T. evansi first | 1 | 31.468 | < 0.0001 |
| | 19:1 same time- 1:19 same time | 1 | 447.546 | < 0.0001 |

Supplementary Table S3 - Full model (A) and *a priori* contrasts (B and C) to test for the impact of initial density and order of arrival on the proportion of females of the two species after two generations of contact separated by type of leaf pair that was added first (B) leaves 2 and 4 first, (C) leaves 3 and 5 first

A) Full model

| **Leaf pair first** | **Factors** | **DF** | **Chisq** | **P-Value** |
|---|---|---|---|---|
| 2-4 | Intercept | 1 | 173.27 | < 0.0001 |
| | Treatment | 6 | 1242.96 | < 0.0001 |
| 3-5 | Intercept | 1 | 178.36 | < 0.0001 |
| | Treatment | 6 | 1094.11 | < 0.0001 |

B) Leaves 2 and 4 first

| Test | Contrasts | DF | Chisq | P-Value |
|---|---|---|---|---|
| Order of arrival | 10:10 same time – 10:10 T. urticae first | 1 | 122.575 | < 0.0001 |
| | 10:10 same time - 10:10 T. evansi first | 1 | 35.093 | < 0.0001 |
| | 19:1 same time- 19:1 T. evansi first | 1 | 0.023 | 0.8795 |
| | 1:19 same time- 1:19 T. urticae first | 1 | 156.773 | < 0.0001 |
| | 10:10 T. evansi first – 10:10 T. urticae first | 1 | 263.084 | < 0.0001 |
| Initial density | 10:10 same time – 19:1 same time | 1 | 95.470 | < 0.0001 |
| | 10:10 same time – 1:19 same time | 1 | 86.782 | < 0.0001 |
| | 10:10 T. urticae first – 1:19 T. urticae first | 1 | 219.233 | < 0.0001 |
| | 10:10 T. evansi first – 19:1 T. evansi first | 1 | 14.213 | < 0.001 |
| | 19:1 same time- 1:19 same time | 1 | 225.344 | < 0.0001 |

C) Leaves 3 and 5 first

| Test | Contrasts | DF | Chisq | P-Value |
|---|---|---|---|---|
| Order of arrival | 10:10 same time – 10:10 T. urticae first | 1 | 4.5076 | 0.03543 |
| | 10:10 same time - 10:10 T. evansi first | 1 | 40.2371 | < 0.0001 |
| | 19:1 same time- 19:1 T. evansi first | 1 | 0.1981 | 0.65628 |
| | 1:19 same time- 1:19 T. urticae first | 1 | 135.319 | < 0.0001 |
| | 10:10 T. evansi first – 10:10 T. urticae first | 1 | 74.0289 | < 0.0001 |
| Initial density | 10:10 same time – 19:1 same time | 1 | 118.394 | < 0.0001 |
| | 10:10 same time – 1:19 same time | 1 | 79.433 | < 0.0001 |
| | 10:10 T. urticae first – 1:19 T. urticae first | 1 | 415.784 | < 0.0001 |
| | 10:10 T. evansi first – 19:1 T. evansi first | 1 | 16.556 | < 0.0001 |

| | | | | |
|---|---|---|---|---|
| | 19:1 same time- 1:19 same time | 1 | 220.497 | < 0.0001 |

Supplementary Table S4 - A) Pairwise contrasts for leaf occupancy in the control treatments (i.e., the single treatments of *T. evansi* and *T. urticae*) after two generations. B) Pairwise contrasts between the leaf occupancy (leaf occupancy/Total females) of the control and each experimental treatment for each leaf. Figures 3 and S3 directly show these differences. All P-values were corrected applying FDR correction. Red indicates significant differences between comparisons.

A)

| **Species** | **Comparison** | **Sum of Squares** | **Df** | **Chisq** | **P-value** |
|---|---|---|---|---|---|
| *T. evansi* | 2-3 | 0.250 | 1 | 306.862 | < 0.000001 |
| | 2-4 | 0.147 | 1 | 836.203 | < 0.000001 |
| | 2-5 | 0.428 | 1 | 17.953 | 0.000136 |
| | 3-4 | 0.341 | 1 | 179.995 | < 0.000001 |
| | 3-5 | 0.691 | 1 | 189.861 | < 0.000001 |
| | 4-5 | 0.813 | 1 | 672.788 | < 0.000001 |
| *T. urticae* | 2-3 | 0.323 | 1 | 56.967 | < 0.000001 |
| | 2-4 | 0.331 | 1 | 51.430 | < 0.000001 |
| | 2-5 | 0.375 | 1 | 25.885 | < 0.000001 |
| | 3-4 | 0.509 | 1 | 0.1564 | 0.99999 |
| | 3-5 | 0.557 | 1 | 6.5275 | 0.06373 |
| | 4-5 | 0.549 | 1 | 4.670 | 0.18417 |

B)

| **Species** | **Comparison** | **Leaf** | **Value** | **Df** | **Chisq** | **P-value)** |
|---|---|---|---|---|---|---|

| | | | | | | |
|---|---|---|---|---|---|---|
| *T. evansi* | 10:10, same time vs control | 2 | 0.43414 | 1 | 12.343 | 0.0014581 |
| | | 3 | 0.59009 | 1 | 35.437 | 3.69E-08 |
| | | 4 | 0.47786 | 1 | 3.5831 | 0.105441 |
| | | 5 | 0.48252 | 1 | 0.9709 | 0.4083083 |
| | 10:10, Tu first vs control | 2 | 0.45123 | 1 | 4.9699 | 0.0515854 |
| | | 3 | 0.47531 | 1 | 2.4914 | 0.1821933 |
| | | 4 | 0.59761 | 1 | 41.7126 | 2.37E-09 |
| | | 5 | 0.38858 | 1 | 37.9931 | 1.14E-08 |
| | 10:10, Te first vs control | 2 | 0.45626 | 1 | 6.3842 | 0.0252857 |
| | | 3 | 0.50572 | 1 | 0.2096 | 0.6968902 |
| | | 4 | 0.52955 | 1 | 7.3861 | 0.015024 |
| | | 5 | 0.45178 | 1 | 9.7311 | 0.0048313 |
| | 19:1, same time vs control | 2 | 0.49922 | 1 | 0.0023 | 0.9790775 |
| | | 3 | 0.5223 | 1 | 3.8566 | 0.0940606 |
| | | 4 | 0.53306 | 1 | 11.4696 | 0.0022009 |
| | | 5 | 0.39777 | 1 | 57.5665 | 1.83E-12 |
| | 19:1, Te first vs control | 2 | 0.42728 | 1 | 23.2653 | 8.99E-06 |
| | | 3 | 0.48824 | 1 | 1.1708 | 0.3679333 |
| | | 4 | 0.54005 | 1 | 17.5827 | 0.0001232 |
| | | 5 | 0.4831 | 1 | 1.4471 | 0.3166341 |
| | 1:19, same time vs control | 2 | 0.53045 | 1 | 0.4926 | 0.5517381 |
| | | 3 | 0.60602 | 1 | 10.3376 | 0.0036497 |
| | | 4 | 0.46321 | 1 | 2.4953 | 0.1821933 |
| | | 5 | 0.43632 | 1 | 3.6407 | 0.1035262 |
| | 1:19, Tu first vs control | 2 | 0.30032 | 1 | 23.1168 | 8.99E-06 |
| | | 3 | 0.55126 | 1 | 1.2563 | 0.3540264 |
| | | 4 | 0.59728 | 1 | 5.5685 | 0.0386425 |
| | | 5 | 0.44459 | 1 | 1.2236 | 0.358196 |
| *T. urticae* | 10:10, same time vs control | 2 | 0.52408 | 1 | 0.8699 | 0.4679865 |
| | | 3 | 0.509 | 1 | 0.1985 | 0.7641754 |
| | | 4 | 0.53742 | 1 | 3.2395 | 0.1388101 |
| | | 5 | 0.44175 | 1 | 8.0739 | 0.0125739 |
| | 10:10, Tu first vs control | 2 | 0.58123 | 1 | 10.6032 | 0.0037188 |
| | | 3 | 0.58773 | 1 | 19.8091 | 6.39E-05 |
| | | 4 | 0.3873 | 1 | 41.0254 | 3.37E-09 |
| | | 5 | 0.53524 | 1 | 2.9614 | 0.1548275 |
| | 10:10, Te first vs control | 2 | 0.48681 | 1 | 0.1913 | 0.7641754 |
| | | 3 | 0.44534 | 1 | 5.675 | 0.0410062 |
| | | 4 | 0.47842 | 1 | 0.812 | 0.4842829 |
| | | 5 | 0.63711 | 1 | 21.4502 | 3.13E-05 |
| | 19:1, same time vs control | 2 | 0.52961 | 1 | 0.1818 | 0.7644475 |
| | | 3 | 0.47676 | 1 | 0.2093 | 0.763149 |
| | | 4 | 0.43147 | 1 | 2.0447 | 0.2443831 |
| | | 5 | 0.62772 | 1 | 3.6032 | 0.1153337 |

| | | | | | | |
|---|---|---|---|---|---|---|
| | 19:1, Te first vs control | 2 | 0.8429 | 1 | 15.9069 | 0.0003105 |
| | | 3 | 0.36771 | 1 | 15.5584 | 0.0003318 |
| | | 4 | 0.52654 | 1 | 0.3927 | 0.6606788 |
| | | 5 | 0.55281 | 1 | 1.2914 | 0.3672987 |
| | 1:19, same time vs control | 2 | 0.43522 | 1 | 6.1247 | 0.033932 |
| | | 3 | 0.54471 | 1 | 3.8059 | 0.105034 |
| | | 4 | 0.47204 | 1 | 1.6515 | 0.2968078 |
| | | 5 | 0.53802 | 1 | 2.4341 | 0.2018333 |
| | 1:19, Tu first vs control | 2 | 0.53274 | 1 | 2.1336 | 0.2373495 |
| | | 3 | 0.49274 | 1 | 0.1735 | 0.7644475 |
| | | 4 | 0.46782 | 1 | 3.4241 | 0.1262504 |
| | | 5 | 0.52949 | 1 | 2.4313 | 0.2018333 |

Supplementary Table S5 - Model to test the impact of type of leaf, treatment and their interaction on female abundance, separated by type of leaf pair that was added first: leaves 2 and 4 first or leaves 3 and 5 first.

| **Leaf pair first** | **Factors** | **DF** | **Chisq** | **P-Value** |
|---|---|---|---|---|
| 2-4 | Intercept | 1 | 31.762 | < 0.0001 |
| | Treatment | 6 | 291.791 | < 0.0001 |
| | Leaf | 3 | 23.005 | < 0.0001 |
| | Treatment*Leaf | 18 | 178.045 | < 0.0001 |
| 3-5 | Intercept | 1 | 19.913 | < 0.0001 |
| | Treatment | 6 | 78.830 | < 0.0001 |
| | Leaf | 3 | 38.996 | < 0.0001 |
| | Treatment*Leaf | 18 | 90.069 | < 0.0001 |

Supplementary Table S6 - Pairwise contrasts for the C-scores of treatments differing in order of arrival or initial density (using the test_interactions function from the phia package). P-values correspond to FDR corrected values. Red indicates significant differences.

| **Comparison** | **Value** | **Df** | **Chisq** | **P-value** |
|---|---|---|---|---|

| | | | | |
|---|---|---|---|---|
| 10:10 same time – 10:10 T. urticae first | -7.004 | 1 | 2.7818 | 0.22246 |
| 19:1 same time- 19:1 T. evansi first | 16.75 | 1 | 0.2975 | 0.73357 |
| 1:19 same time- 1:19 T. urticae first | 25.615 | 1 | 0.1471 | 0.77514 |
| 10:10 T. evansi first – 10:10 T. urticae first | -3.895 | 1 | 6.7736 | 0.06476 |
| 10:10 same time – 10:10 T. evansi first | -9.165 | 1 | 1.6914 | 0.31244 |
| 10:10 same time – 19:1 same time | -3.16 | 1 | 10.3562 | 0.02175 |
| 10:10 same time – 1:19 same time | -6.551 | 1 | 3.3073 | 0.22246 |
| 10:10 T. urticae first – 1:19 T. urticae first | -7.427 | 1 | 2.261 | 0.25328 |
| 10:10 T. evansi first – 19:1 T. evansi first | -5.942 | 1 | 3.6646 | 0.22246 |
| 19:1 same time- 1:19 same time | 6.104 | 1 | 2.8884 | 0.22246 |

Supplementary Table S7 - A priori contrasts for the expression of gene known to respond to spider mite infestation (using the test_interactions function from the phia package). PI-IIc and PPOD genes are from the jasmonic acid pathway and PR1a from the salicylic acid pathway. Santpoort and Viçosa populations correspond to the positive (induction) and negative (suppression) controls. Red indicates significant differences.

| Comparison | Gene | Value | Df | Chisq | P-value |
|---|---|---|---|---|---|
| Santpoort vs *T. urticae* outbred | PI-IIc | 40.926 | 1 | 32.5031 | 1.190e-08 |
| | PPOD | 5.1702 | 1 | 5.7057 | 0.01691 |
| | PR-1a | 3.9201 | 1 | 8.0322 | 0.004595 |
| Santpoort vs *T. evansi* outbred | PI-IIc | 31.662 | 1 | 28.1636 | 1.115e-07 |
| | PPOD | 3.2132 | 1 | 2.8802 | 0.08968 |
| | PR-1a | 4.1788 | 1 | 8.8014 | 0.003010 |
| Viçosa vs *T. urticae* outbred | PI-IIc | 1.141 | 1 | 0.1554 | 0.6934 |
| | PPOD | 0.8993 | 1 | 0.0238 | 0.87736 |
| | PR-1a | 1.6909 | 1 | 1.1875 | 0.275841 |
| Viçosa vs *T. evansi* outbred | PI-IIc | 1.475 | 1 | 0.3560 | 0.5507 |
| | PPOD | 1.4470 | 1 | 0.2886 | 0.59109 |
| | PR-1a | 1.5862 | 1 | 0.9161 | 0.338511 |

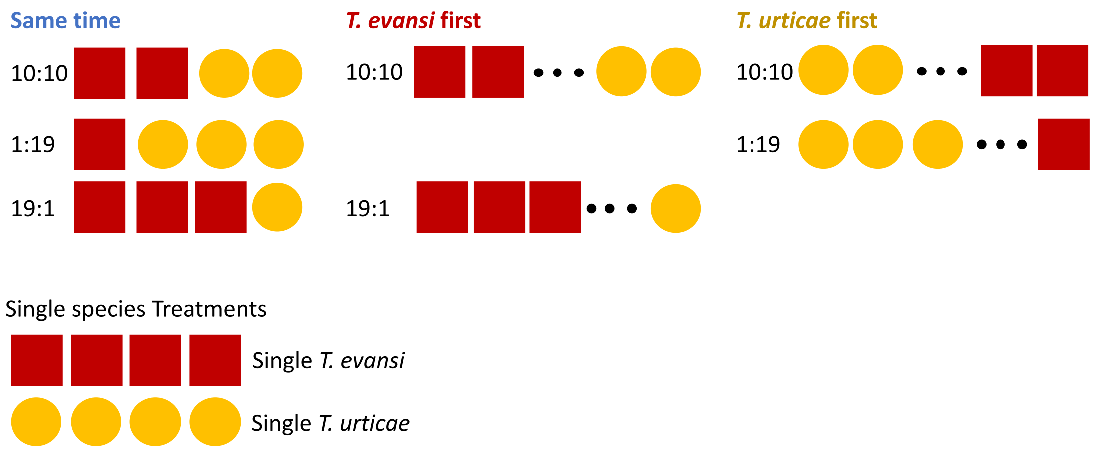


Figure S1 – Experimental setup used to test the impact of order of arrival and initial frequency on competition between *T. urticae* and *T. evansi*. Red squares code for *T. evansi*, and yellow circles code for *T. urticae*.

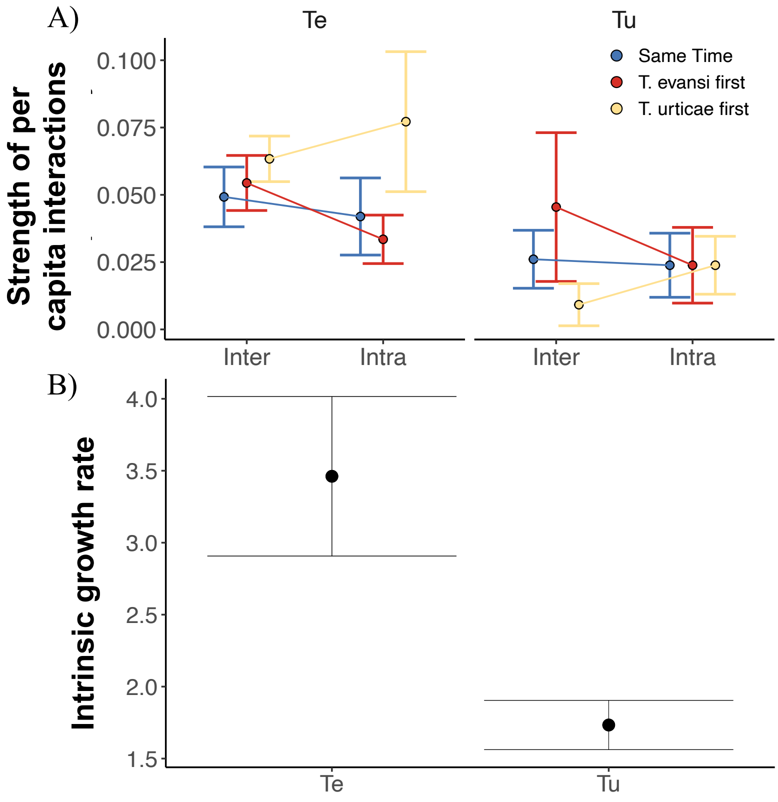


Figure S2 – Changes in the strength of per-capita interactions (A) and the intrinsic growth rate in absence of competitors (lambda in model 6, see main text) (B) for *T. urticae* and *T. evansi*

when they arrive at the same time (yellow), *T. evansi* arrives first (red) and *T. urticae* arrives first (blue). Error bars represent 95% confidence intervals of the estimates.

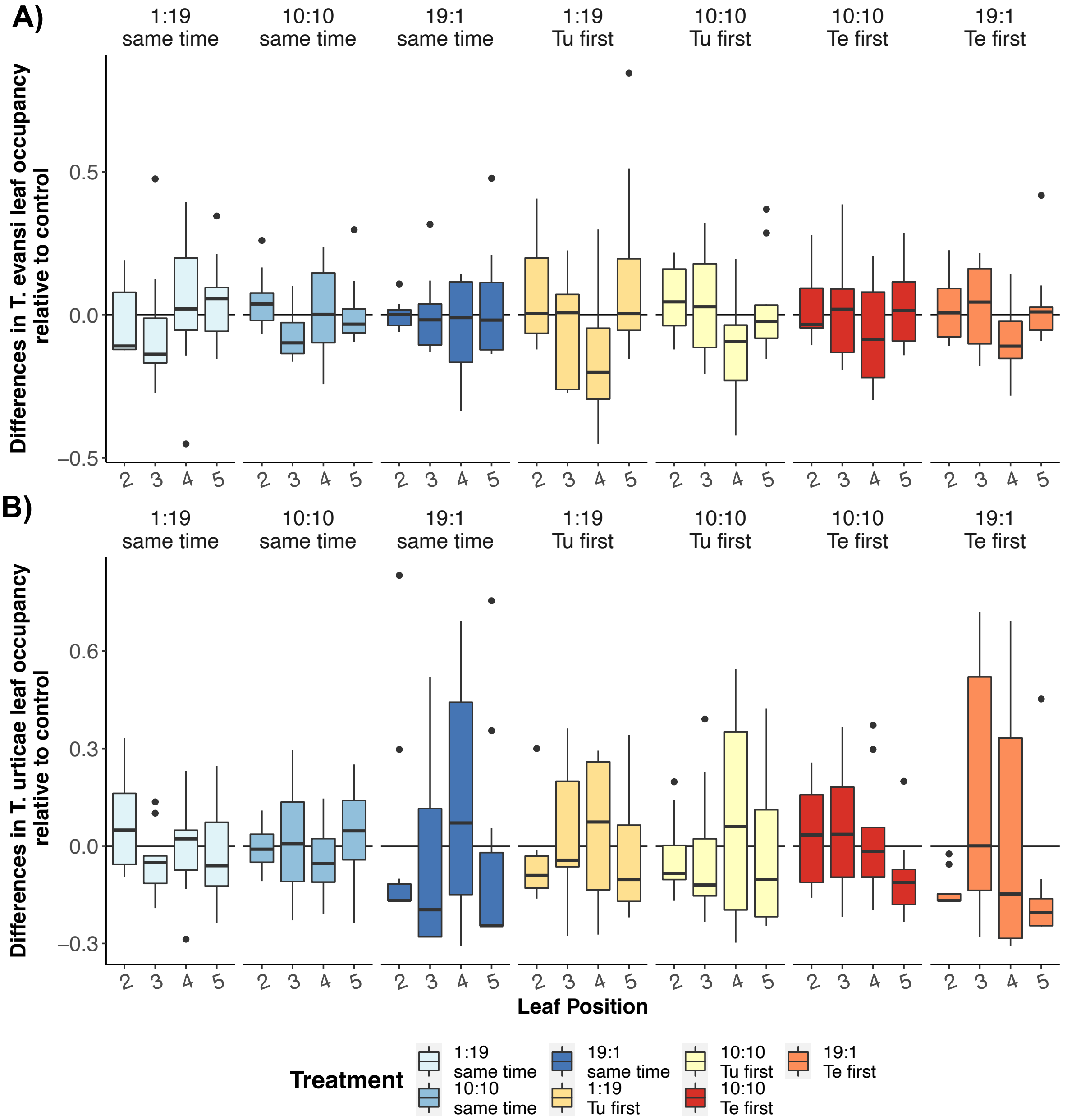


Figure S3– Difference in *T. evansi* (A) and *T. urticae* (B) occupancy relative to the control (y-axis) across leaves (x-axis) for the different experimental treatments. For each species in each treatment, we first calculated the proportion of females occupying that leaf in relation to the total number of females in each box. The we computed the difference between the proportion on each leaf/box and the average proportion of females occupying that leaf in the single species treatment (control). Thus, positive values indicate that, in that treatment, more females are present in that leaf/side than when the species is by itself and negative values indicate the reverse pattern.

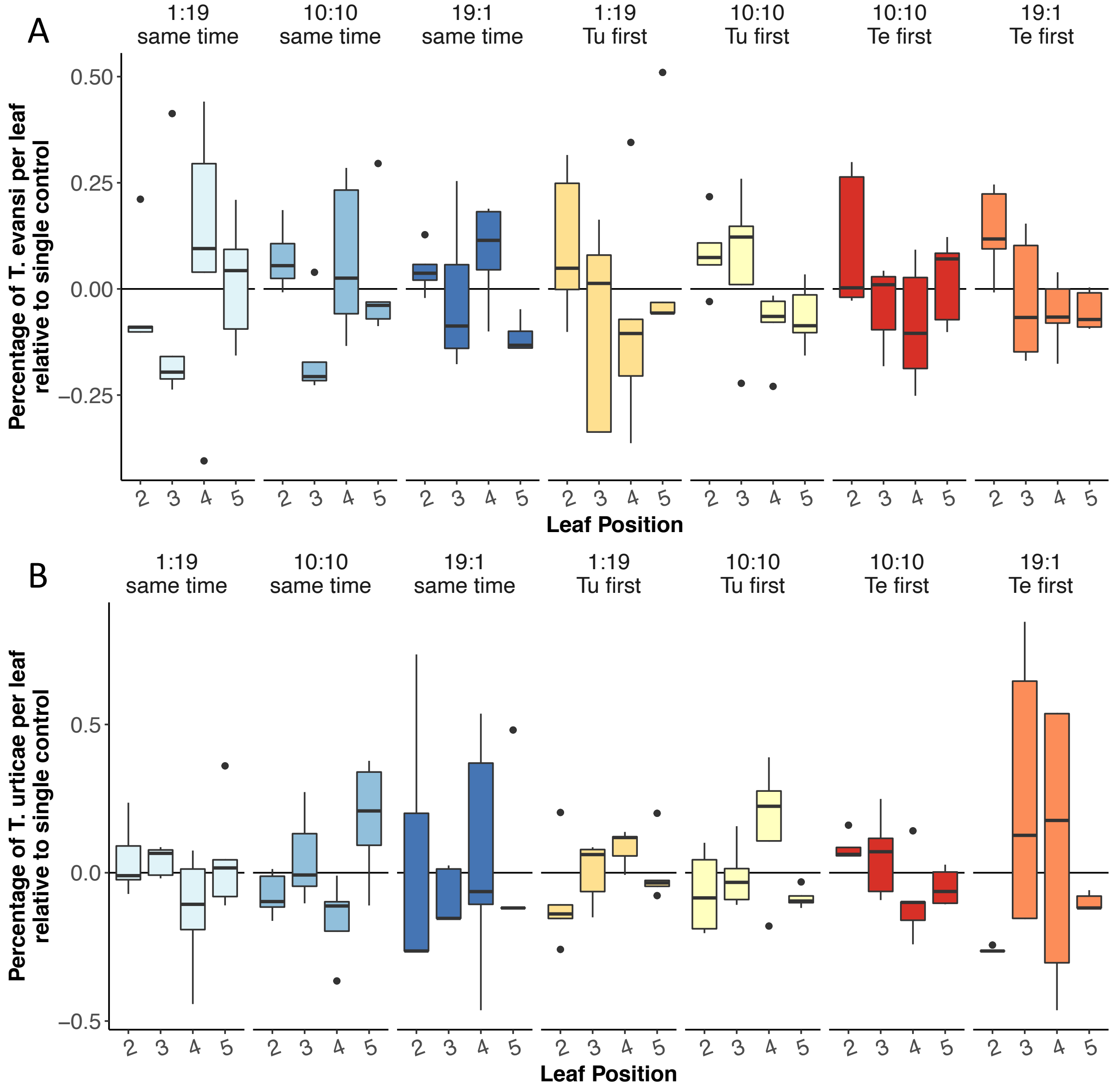


Figure S4– Difference in *T. evansi* (A) and *T. urticae* (B) occupancy relative to the control (y-axis) across leaves (x-axis) when accounting for leaf order. For each species in each treatment, we first calculated the proportion of females occupying that leaf in relation to the total number of females in each box separately for each leaf order (either 2-4 first or 3-5 first). Then we computed the difference between the proportion on each leaf/box and the average proportion of females occupying that leaf in the single species treatment (control) separated by leaf order. The plot was done using data from the two types of leaf order. The overall pattern is similar to that obtained in the data without accounting for the changes in leaf order.

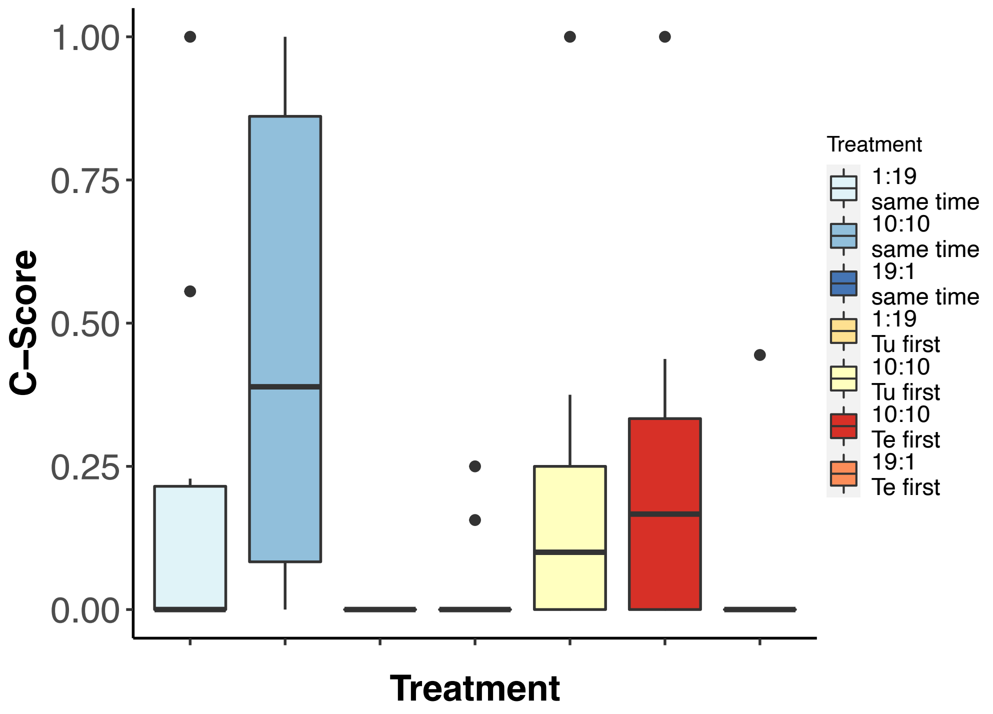


Figure S5 – C-score (y-axis) for the different experimental treatments (x -axis). The score varies between 0 (no aggregation) and 1 (aggregation).

A)

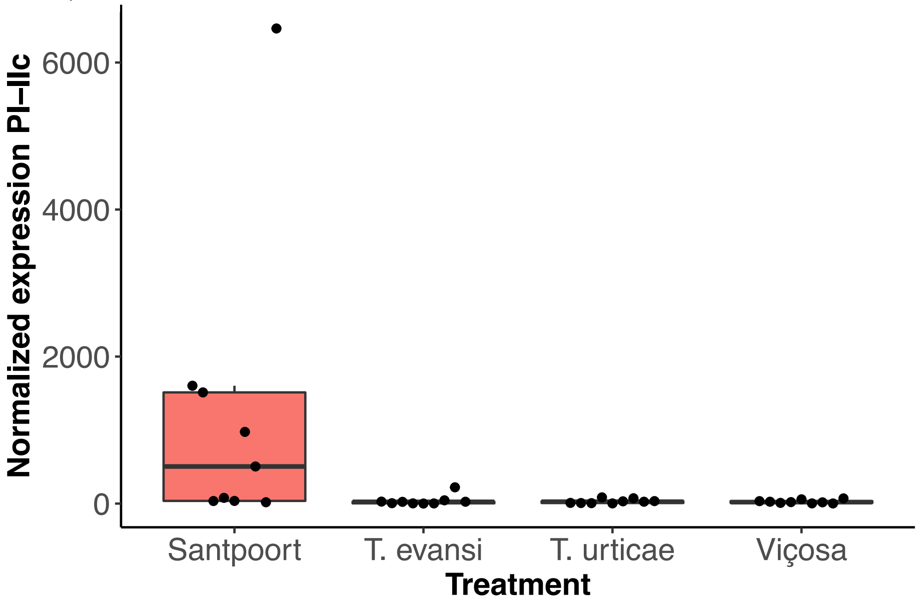
Normalized expression PI–IIc
6000
4000
2000
0
Santpoort
T. evansi
T. urticae
Viçosa
Treatment

B)

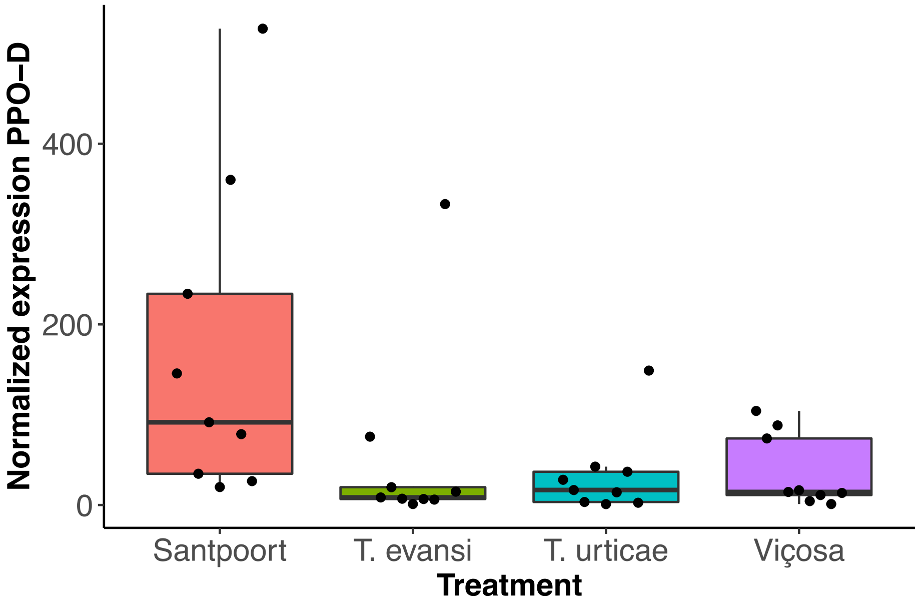
Normalized expression PPO–D
400
200
0
Santpoort
T. evansi
T. urticae
Viçosa
Treatment

C)

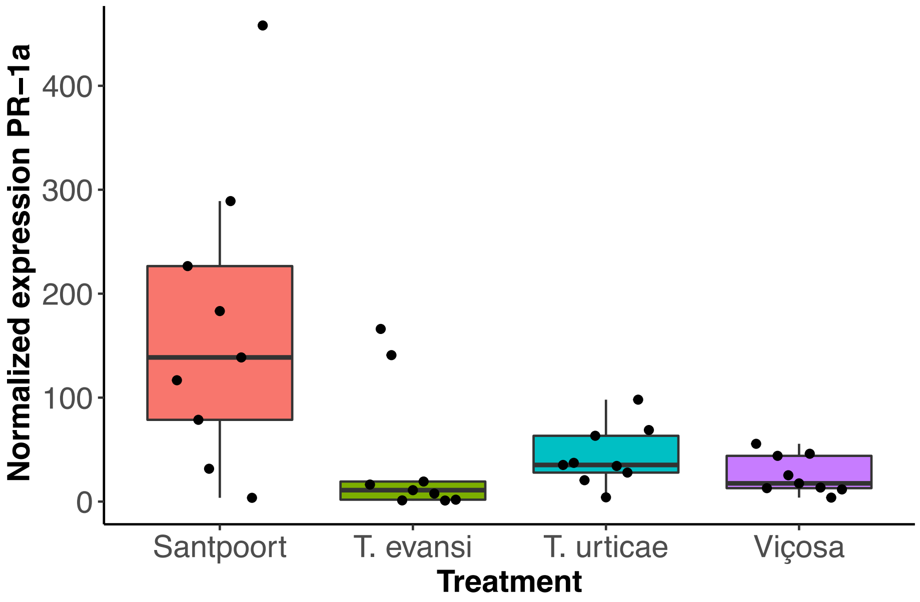


Figure S6 – Normalized gene expression of PI-IIc (A), PPOD (B) and PR-1a (C) genes. Santpoort and Viçosa are control populations for induction and suppression, respectively. Santpoort populations induce gene expression for all three genes, whereas the other three populations show low expression of all genes in general.
